# Hidden palette: Culturing the sauropsid gut microbiome reveals a high prevalence and diversity of bacteria that undergo carotenoid biosynthesis

**DOI:** 10.64898/2026.03.06.710123

**Authors:** Alix E. Matthews, Victoria Gates, Kartikey Sharma, Sebastian Gallego, Madeline Desmond, Brian Trevelline, Marcella D. Baiz

**Author notes:** Corresponding author:* Marcella D. Baiz; 135 Founders Promenade, Cooke Hall 109, Buffalo, NY 14260-1300.

## Abstract

Pigments play essential roles in communication, sexual signaling, and ecological adaptation in taxa across the tree of life and are especially important for animals, which often rely on carotenoid pigments for their coloration. One potentially important yet largely overlooked source of pigments is host-associated microbes. Microbial carotenoids may influence host coloration or other fitness-related traits, but the extent to which they occur or vary across associations with host species remains largely unknown. To begin to address this gap, we used culture-based techniques to isolate gut-associated bacteria from multiple wild sauropsid species. We cultured 157 morphologically distinct bacterial isolates from 25 individuals representing eight host species; 17% of isolates were yellow- or orange-pigmented, and 40% of isolates exhibited absorbance spectra consistent with the presence of carotenoids. Sequencing of the 16S rRNA gene revealed that isolates spanned four phyla and 35 genera. Isolates that were consistent with the presence of carotenoids were found across all four phyla, with *Pantoea* being the most frequently represented carotenoid-associated genus. These results demonstrate that carotenoid-capable bacteria are both taxonomically diverse and prevalent in the gut microbiomes of wild sauropsids. More broadly, our findings provide insight into a hidden dimension of host-microbe interactions and their potential function.

## INTRODUCTION

Natural pigments underlie functionally important phenotypes in taxa across the tree of life. For example, chlorophyll, a green pigment found in plants, cyanobacteria, and algae, is essential for initiating photosynthesis. The colorful plumage of many bird species is also produced by pigments such as carotenoids, melanins, and porphyrins (among others); these pigment-based plumages are critical for sexual signaling and other types of communication (Hill 1991; Hedley and Caro 2022). Microbes express colorful pigment-based phenotypes which are functionally important and can sometimes be very salient; for instance, the pinkish color of hypersaline lakes is attributed to carotenoid pigments synthesized by halophilic microbes in the water, where these molecules play key roles in photoprotection and antioxidant defense (Teller 1987; Mandelli *et al*. 2012). However, unless grown in culture, the color phenotypes of most microbes are usually inconspicuous. This is especially true for host-associated communities, where microbes are dispersed within complex ecological communities and remain hidden from view. While many microbes in such diverse communities may still express pigments, their presence is often overlooked, and their proximate and ultimate functions remain poorly understood. In particular, the ecological and functional roles of microbial pigments in host-microbe interactions are largely unexplored, despite their potential to mediate important host phenotypic traits, such as those involved in communication and sexual signaling.

Unlike bacteria, most animals cannot synthesize carotenoid pigments *de novo* and must obtain them from external sources (Toews, Hofmeister and Taylor 2017). Carotenoid acquisition has therefore traditionally been assumed to occur from diet alone (Hill 1992), including diets supplemented with microbially produced carotenoids in commercial feeds (Pasarin and Rovinaru 2018; Al-Monofy *et al*. 2025). Recent research highlights a potential link between host-associated carotenoid-synthesizing microbes and host phenotypes. For example, in both invertebrate (e.g., amphipods [Koellsch, Poulin and Salloum 2024], molluscs [Liu *et al*. 2020]) and vertebrate hosts (e.g., fish [Nguyen *et al*. 2020], birds [Baiz, Wood and Toews 2024]), correlations have been observed between the presence of carotenoid-producing bacterial taxa and variation in host coloration. In birds specifically, the abundance of certain bacterial taxa in the gut microbiome predicted to encode carotenoid biosynthesis genes (e.g., *Sphingomonas*) is positively associated with the extent of carotenoid-based yellow plumage color of their hosts, suggesting a potential microbial contribution to host coloration (Baiz, Wood and Toews 2024). Beyond coloration, carotenoids also support host immune function (Blount *et al*. 2003) and act as antioxidants (Stahl and Sies 2003). Due to trade-offs in the allocation of carotenoids to signal phenotypes versus maintenance, carotenoid-based coloration is widely thought to be an honest indicator of mate quality (Hill 1991; Freeman-Gallant *et al*. 2011). Whether (and to what extent) bacterial carotenoids contribute to host coloration, immunity, or reproductive success remains unknown, leaving a key gap in our understanding of the microbiome’s role in host fitness and sexual selection. Addressing this gap requires characterizing the diversity and estimating the prevalence of host-associated microbes that are capable of synthesizing carotenoids.

Bacterial taxa across at least 12 phyla are known to synthesize carotenoids, producing more than 300 distinct compounds (Yabuzaki 2017). Some well-characterized carotenoid producers include human pathogens such as *Staphylococcus aureus*, which produces staphyloxanthin (Pelz *et al*. 2005) and draws its specific epithet from its golden color (Rosenbach 1884). They fill various ecological roles and are found in diverse environments, from marine ecosystems (Du *et al*. 2006) to extreme Antarctic habitats (Dieser, Greenwood and Foreman 2010), as well as host-associated microbiomes (Yoon *et al*. 2011; Kim *et al*. 2015). Several bacterial taxa commonly found in host-associated microbiomes are predicted to synthesize carotenoids, including *Pseudomonas*, *Rhizobium*, and *Sphingomonas* (Escallón, Belden and Moore 2019; Baiz, Wood and Toews 2024). Given the broad taxonomic and ecological distribution of carotenoid-producing bacteria and the complexity of their biosynthetic pathways, carotenoids may be widely advantageous for bacteria to synthesize. Indeed, carotenoids prevent photodamage and oxidative stress (López *et al*. 2023), protect cells from extreme environmental stressors (Dieser, Greenwood and Foreman 2010), and even act as virulence factors (Liu *et al*. 2005). However, the roles and possible advantages of carotenoid biosynthesis by symbiotic bacteria in the host gut microbiome remain uncharacterized. Evaluating the extent and diversity of carotenoids synthesized by gut symbionts is a critical first step to understand how microbial carotenoids may influence host fitness and the stability of host-microbe symbioses.

Here, we isolated phenotypically unique bacteria from the gut microbiomes of eight sauropsid host species to study their pigment phenotypes and genotypic diversity. We mainly focused our sampling on colorful wood warblers (family Parulidae), though we also included two avian species which do not express carotenoid-pigmented plumage as well as one lizard species to broaden the evolutionary representation of hosts. Our primary objectives were to (1) characterize the phenotypes and genotypes of gut bacteria across host species, and (2) test the hypothesis that the host gut microbiome includes bacteria capable of carotenoid biosynthesis. We examined bacterial diversity by reconstructing the phylogenetic history of our isolates and further tested whether they exhibit host- or environment-specific distributions.

## METHODS

### Fieldwork

We collected fecal samples from 20 wild birds between 03-27 June 2024 across five sites in western New York State, USA. Avian hosts included five species within the family Parulidae: Blue-winged Warbler (BWWA; *Vermivora cyanoptera*), Common Yellowthroat (COYE; *Geothlypis trichas*), Chestnut-sided Warbler (CSWA; *Setophaga pensylvanica*), Hooded Warbler (HOWA; *Setophaga citrina*), and Yellow Warbler (YEWA; *Setophaga petechia*), one species within the family Mimidae: Gray Catbird (GRCA; *Dumetella carolinensis*); and one species within the family Hirundinidae: Purple Martin (PUMA; *Progne subis*). We captured individuals in a mist net (parulids and Gray Catbirds) or at the nest box (Purple Martins) and immediately placed them in a sterile sample collection chamber made of a paper bag lined with aluminum foil which sat underneath a ¼ inch steel mesh platform separating the bird from the fallen fecal material. After ten minutes, we removed the bird from the chamber and collected whole feces by using sterile inoculating loop to transfer it into a tube containing 750µL of 30% glycerol in PBS. After returning to the lab, we vortexed the sample tubes and transferred half of the volume to a second tube and froze both tubes at -80°C until they were used for bacterial cultivation. We also collected a field negative control by sampling an empty sterile collection chamber in the same manner as described above.

We captured adult wild common wall lizards (POMU; *Podarcis muralis*) on 06 June 2024 from urban, roadside stone walls in Cincinnati, Ohio, USA using a thread lasso attached to an extendable fishing rod. We placed each lizard into individual nylon, breathable cloth sacks and then placed all sacks into a cooler for transport to the laboratory at Kent State University. In the laboratory, we housed lizards in individual plastic rat cages (18” L x 9.25” W x 6.5” H) with a mesh lid. We used a 40W incandescent heat bulb suspended above one end of the cage to maintain a temperature gradient from ∼25-40°C within cages, allowing for natural behavioral thermoregulation. Additionally, we placed a 39W 10.0 UVB light tube horizontally across cages to provide a source of UV light. UV and ambient light were provided on a 14hr:10hr daily light:dark cycle with heat bulbs turned on in alternating one hour on/one hour off intervals for a total of six hours per day during the diurnal light cycle. Inside each cage, lizards were provided with a basking platform doubling as a shelter and a Petri dish filled with distilled water. We replaced the water and misted cages daily. We fed lizards every other day a diet of crickets and mealworms dusted with reptile vitamin supplement *ad libitum*. Between 14-16 June 2024, 2-3 fecal samples were opportunistically collected from three lizards. The three lizards were selected because they represent three distinct common wall lizard color morphs (orange, yellow, and white; Andrade *et al*. 2019). Individual fecal samples were removed from the cage using sterilized forceps and placed in sterile cryovials containing 1mL of sterilized 10% glycerol in PBS. The samples were stored at -80°C until bacterial cultivation.

### Bacterial Cultivation

Fecal samples and the negative field control were vigorously vortexed and serially diluted from 10^−1^ to 10^−6^. For aerobic cultivation of avian fecal samples, we used Brain Heart Infusion (BHI; non-selective) medium for cultivation of generalist taxa and “L9” (Yim *et al*. 2010) medium for selective cultivation of *Sphingomonas* spp. For aerobic cultivation of lizard fecal samples, we used BHI and Tryptic Soy Agar (TSA; non-selective). For each dilution, we prepared two replicate agar plates (100 x 15mm) under a biosafety cabinet. On each plate, we spread 100µL of diluted sample using 5-10 sterile glass beads (Zymo Cat. S1001). We included a negative (30% glycerol in PBS) and positive (*Sphingomonas paucimobilis*; ATCC 10829) control for each batch of samples processed. Plates were incubated aerobically at 30°C (lizard samples) or 35°C (avian samples) for 24-48 hours or until sufficient growth occurred. We selected total cultures exhibiting distinct, well-isolated colonies of bacteria for phenotypic characterization of color, form, margin shape, texture, size, opacity, and elevation (Table S1). Colonies demonstrating distinct morphologies were then isolated by streaking on fresh agar plates. Once inoculated, we incubated the plates at 30°C or 35°C and monitored them every 24 hours to prevent overgrowth.

For anaerobic cultivation (avian samples only), BHI plates were prepared and pre-reduced by placing them in an anaerobic container (Mitsubishi Gas Chemical) overnight. Samples were processed as previously described, but our positive control was *Fusobacterium nucleatum* subsp. *polymorphum* (ATCC 10953). Plates were immediately placed in anaerobic containers along with AnaeroPacks (Mitsubishi Gas Chemical) and an anaerobic indicator to create and confirm an oxygen-free environment. Containers were incubated at 35°C for 48-72 hours. After incubation, unique colonies were phenotypically characterized, selected, and streaked onto fresh plates as previously described, but under anaerobic conditions.

We collected individual colony isolates from T-streak plates that were approximately 0.5cm^2^ in size with a sterile loop and incubated them in 1.5-2mL of sterile broth to grow liquid cultures. We placed the samples (including a negative control containing only sterile broth) into an orbital incubator at 35°C at 120rpm for 18-24 hours under either aerobic or anaerobic conditions. Half of the liquid culture was then mixed in 30% glycerol in PBS for long-term storage and the remaining half was centrifuged and decanted such that the pellets could be used as input for PCR and sequencing (below).

### 16S Amplification and Sequencing

For each isolate cultured from avian samples, we amplified the full-length 16S rRNA gene (including variable regions V1 to V9) by performing colony PCR amplification using 5µM universal 16S primers 27F (5’-AGAGTTTGATCMTGGCTCAG-3’) and 1492R (5’-CGGTTACCTTGTTACGACTT-3’; Lane 1991). PCRs consisted of 6.5µL nuclease-free water, 2µL forward primer, 2µL reverse primer, 12.5µL LongAmp Hot Start*Taq* 2X Master Mix (New England BioLabs, M0533S), and approximately 0.2µL of pelleted bacterial cells (i.e., gently touching a pipette tip to the pellet). Reaction conditions were as follows: initial denaturation at 94°C for 30s, followed by 30 cycles of 94°C for 30s, 55°C for 40s, 65°C for 90s, followed by a final extension at 65°C for 10 min. PCR products were visualized on 1% agarose gels with SYBR Safe gel stain. Isolates that failed to amplify using colony PCR were regrown on plates and DNA was extracted with the ZymoBIOMICS DNA Miniprep Kit (D4300). PCRs were repeated under the same conditions using 2µL of extracted DNA. We purified PCR products using Serapure beads (0.6x volume) and eluted in 25µL of 10µM Tris-HCl. The cleaned products were quantified with a Qubit 3.0 Fluorometer (Invitrogen) using the dsDNA HS Assay Kit (Q32854).

To prepare sequencing libraries, we used the Rapid Sequencing V14 Barcoding Kit (Oxford Nanopore Technologies [ONT], SQK-RBK114.98) following the manufacturer’s instructions (version RAA_9198_v114_revK_11Dec2024) with slight modifications. Specifically, we prepared half-size reactions (e.g., 25ng of amplicon DNA in 4.5µL of volume instead of 50ng in 9µL of volume) and doubled the incubation times (e.g., 20 min instead of 10 min). The barcoded and pooled libraries were loaded on a Flongle Flow Cell (ONT, FLO-FLG114) that was fitted to a MinION Mk1B device (ONT, MIN-101B) and sequenced. We used MinKNOW software (version 24.11.10) for raw data collection. We specified high accuracy basecalling and filtered reads to a minimum quality score of 9 and a minimum read length of 200bp. We used the EPI2ME (desktop application version 5.2.3) “wf-amplicon” workflow (version 1.1.4) to trim adapters and generate *de novo* consensus sequences for each barcoded amplicon. We downsampled to 500 reads (minimum of 150) and specified a maximum read length of 2000bp. Raw reads are deposited in the NCBI Short Read Archive (SRA) database (Table S2).

Samples that failed to meet these thresholds or failed during consensus sequence generation (n = 5) were alternatively sequenced (Sanger) in both directions by the Roswell Park Comprehensive Cancer Center DNA Sequencing Lab on an Applied Biosystems 3500 Genetic Analyzer (avian samples). All lizard samples (n = 5) were sequenced (Sanger) via direct colony sequencing on an Applied Biosystems 3730xl DNA Analyzer at Genewiz (Azenta). Forward and reverse sequences were assembled using the “isolateR” package (Daisley et al. 2024), specifying a sliding window cutoff of 20. Sanger sequences are deposited in GenBank (Table S2).

### Taxonomic Classification, Phylogenetic, and Diversity Analyses

We uploaded our consensus sequences to the Alignment, Classification and Tree Service on the SILVA website (https://www.arb-silva.de/aligner/), which uses SINA version 1.2.12 (Pruesse, Peplies and Glöckner 2012). We enabled the “search and classify” option to classify our sequences against the SILVA database; we specified a minimum identity of 95% and rejected sequences with <70% identity (Quast *et al*. 2013). For isolates that were not identified to the genus level with SILVA, we used the BLASTn program on the NCBI website (https://blast.ncbi.nlm.nih.gov/; Altschul *et al*. 1990) and classified the genus based on the most common genus of the top 50 hits as sorted by e-value.

We ensured all reads were oriented in the same direction before combining into a single FASTA for alignment. We performed multisequence alignment with MAFFT version 7.525 (Katoh and Standley 2013) using the default parameters (--auto). We included *Adhaeribacter pallidiroseus* (phylum Bacteroidota; NCBI accession NR_181905) as our outgroup to root the tree due to its ancestral placement from the ingroup taxa we identified. We estimated a maximum likelihood phylogeny using the alignment (1,541bp) in IQ-Tree version 2.4.0 (Minh *et al*. 2020). We used 1000 ultrafast bootstrap replicates and 1000 SH-aLRT replicates as measures of branch support. The best model of nucleotide substitution was estimated in IQ-Tree prior to the analysis using ModelFinder (Kalyaanamoorthy *et al*. 2017) and corrected Akaike Information Criterion (AICc).

To define amplicon sequence variants (ASVs), we first pre-clustered sequences into operational taxonomic units (OTUs) at 99% identity using VSEARCH version 2.30.2 (“cluster_fast”), with minor manual adjustments based on monophyly. Then, sequences within each OTU were aligned using MAFFT, and leading and trailing gaps were removed to define a shared homologous region. Trimmed sequences were then dereplicated using VSEARCH (“fastx_uniques”) into unique ASVs.

We tested whether microbiome community composition was structured by host species identity or sampling site using PERMANOVA. We generated a matrix of ASV presence/absence among samples and calculated unweighted UniFrac distance using the “distance” function in phyloseq version 1.50.0 (McMurdie and Holmes 2013). Unweighted UniFrac distance estimates phylogenetic dissimilarity among communities based on branch lengths in the ASV phylogeny. We conducted PERMANOVA tests on the unweighted UniFrac distance matrix using the “adonis2” function in vegan version 2.6-10 (Oksanen *et al*. 2025) by assessing the marginal effects of each term (by=‘margin’). We also tested for differences in alpha diversity between species and sites by estimating Faith’s phylogenetic diversity (PD) in each sample using the “estimate_pd” function in btools version 0.0.1 (Battaglia 2019), fitting a linear model and running a Type II analysis-of-variance (ANOVA) using car version 3.1-3 (Fox and Weisberg 2019).

### Analysis of Carotenoids

To assess whether bacterial pigments fell within the expected range of wavelengths at maximum absorbance for carotenoids, we performed an absorbance assay. To do so, we re-cultured isolates from stocks in sterile broth under either aerobic or anaerobic conditions to maximum density based on OD600 values. Samples were centrifuged into pellets and the supernatant was decanted. We added 800µL of a methanol:acetone (7:3 V/V) solution to extract pigments from the bacterial pellets (Toomey *et al*. 2022). Samples were then either directly incubated at 60°C for 90 min at 250rpm on a shaking incubator (avian samples) or were first sonicated using a sonicating wand for 20 seconds at 22,000 Hz then incubated at 60°C for 90 min at 400rpm on a shaking incubator (lizard samples). A negative control containing only the methanol:acetone solution was also included. Following incubation (and once cells were visibly devoid of color), we centrifuged samples to pellet any cell debris. We then transferred 200µL of the supernatant into a 96-well plate, including the negative control. We performed absorbance assays on a Tecan Spark multimode plate reader using the SparkControl program version 2.3 (avian samples) and on a Biotek Synergy H1 multimode plate reader using the Gen5 program version 1.11 (lizard samples). We measured absorbance from 300-700nm (avian samples) or from 260-700nm (lizard samples), taking a reading every 2nm. Absorbance values were standardized relative to the negative control in each plate.

We considered absorbance spectra to be consistent with a carotenoid if the wavelength at maximum absorbance (lambda max) fell between 378nm-512nm (Rodriguez-Amaya 2001). To identify lambda max, we used the function “find_peaks” in the R package photobiology version 0.14.0 (Aphalo 2015) to detect lambda values larger than five consecutive values on either side (span = 11, global.threshold = 0.5). Three researchers (SG, VG, KS) independently inspected the spectra for validation (Figure S1). If two or more observers disagreed with the results of “find_peaks” for a certain sample, we considered that spectrum as “missing data” as it likely represents either a false positive or false negative.

## RESULTS

### Phenotypic diversity

We cultured a total of 152 morphologically distinct bacterial isolates across 20 individual birds and five distinct isolates across three individual lizards (total n = 157; Table 1). *Sphingomonas paucimobilis* grew on our selective (“L9”) media positive control in aerobic conditions and *Fusobacterium nucleatum* subsp. *polymorphum* grew on our non-selective media in anaerobic conditions. No bacteria grew on our negative field control in either aerobic or anaerobic conditions.

**Table 1.** Number of phenotypically unique isolates by host species (species code in parentheses) and media type. N represents the number of host individuals.

| Host Species | N | Aerobic |  | Anaerobic |
| --- | --- | --- | --- | --- |
|  |  | Non-selective | Selective | Non-selective |
| Purple Martin (PUMA) | 4 | 24 | 3 | 12 |
| Blue-winged Warbler (BWWA) | 5 | 22 | 0 | 9 |
| Common Yellowthroat (COYE) | 3 | 11 | 0 | 6 |
| Chestnut-sided Warbler (CSWA) | 3 | 14 | 1 | 9 |
| Gray Catbird (GRCA) | 2 | 8 | 3 | 5 |
| Hooded Warbler (HOWA) | 1 | 6 | 1 | 4 |
| Yellow Warbler (YEWA) | 2 | 10 | 0 | 4 |
| Common Wall Lizard (POMU) | 3 | 5 | NA | NA |

We characterized initial colony phenotype upon growing total cultures, but herein we report only their phenotype in isolation (i.e., after T-streaking), as we occasionally observed minor differences. Across both anaerobic and aerobic conditions and both non-selective and selective media, 79% of isolates were shades of white or cream (Table S1). This pattern was generally consistent across oxygen conditions and media as well, such that ivory white was the most frequent color for aerobic non-selective media, pure white was the most frequent color for aerobic selective media, and off-white cream was the most frequent color for anaerobic non-selective media (Figure 1). We observed four distinct shades of yellow and one shade of orange across isolates (Table S1; Figure 1)

**Figure 1.**
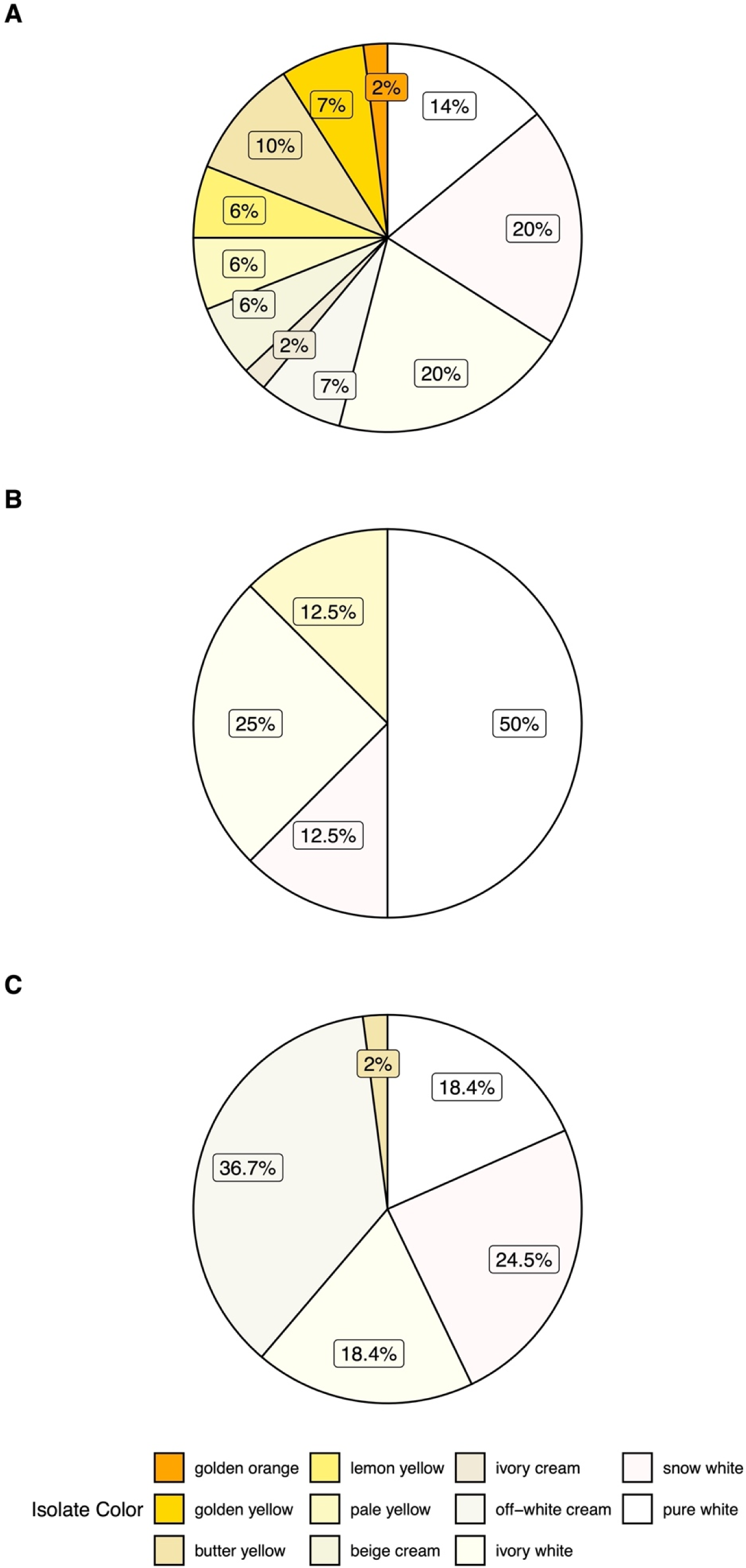
Distribution of color phenotypes observed in 157 isolates cultured from 20 individual birds and three lizards on (A) non-selective aerobic, (B) non-selective anaerobic, and (C) selective aerobic media.

We further characterized isolates based on their overall shape (“form”), relative size, edge (“margin”) texture, opacity, and elevation (relative height). The most common shape of isolates was circular (75.9% of aerobic colonies, 85.7% of anaerobic colonies), with the majority having a surface texture being described as moist and shiny (84.3% of aerobic colonies, 85.7% of anaerobic colonies). The variation in size was more evenly distributed among aerobic colonies than anaerobic isolates: 27.7% aerobic isolates were described as tiny, 32.4% were described as small, 30.5% as medium, whereas 77.6% of anaerobic isolates were described as small. The edge texture of colonies was primarily smooth (79.6% of aerobic colonies, 73.5% of anaerobic colonies). Most isolates grown in aerobic conditions were either opaque (53.7%) or translucent (40.7%), and a similar pattern was observed for anaerobic isolates (38.8% opaque; 42.9% translucent). Most isolates were flat (69.4% of aerobic, 55.1% of anaerobic), but many were also described as umbonate (i.e., having a slightly raised bump in the center of the colony; 13.9% of aerobic, 38.8% of anaerobic). All phenotype data can be found in Table S2.

We performed absorbance assays on 156 of the 157 morphologically distinct bacterial isolates (one isolate failed to regrow for this assay). Lambda max values fell within the expected range for carotenoids for 62 isolates (39.7%). These isolates belong to 19 genera, with *Pantoea* and *Enterococcus* most highly represented (Figure 2; Table S2). All eleven streak colors were represented among these isolates, with 29 isolates (46.8%) being a shade of yellow or orange. The majority of these 62 isolates were cultivated under aerobic conditions (82.3%). There were 74 isolates (47.4%) with lambda max values that fell outside of the expected carotenoid range, representing 21 genera. *Lactococcus* and *Enterococcus* were the two most prevalent genera in this group. Streak colors were generally shades of white and cream, although two lemon yellow isolates and one pale yellow isolate were included with this group. The cultivation conditions were more evenly split for isolates whose lambda max values fell outside of the expected range for carotenoids (58.1% aerobic, 41.9% anaerobic). We could not determine lambda max for 20 isolates (12.8%).

**Figure 2.**
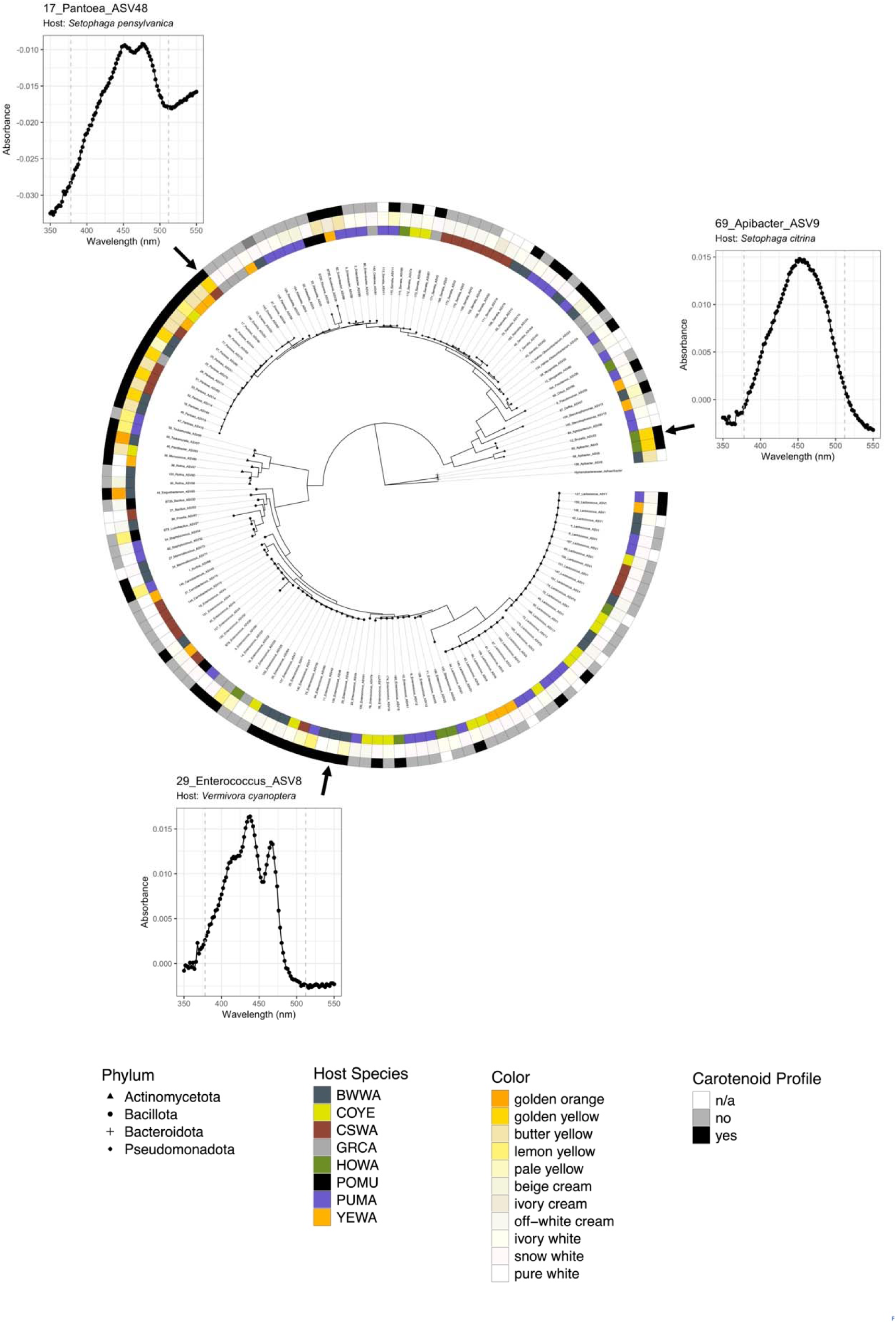
Maximum likelihood phylogenetic tree estimated from the 16S rRNA gene of 147 bacterial isolates cultured from wild sauropsid fecal samples. Tips are labeled with the genus and ASV of each isolate, and shapes at the end of tips represent their taxonomic classification at the level of phylum. The innermost ring represents the host species from which the isolate was isolated; the middle ring represents the color of the isolate; the outermost ring represents whether the isolate is consistent (“yes”) or inconsistent (“no”) with a carotenoid based on absorbance spectra (tips with “n/a” could not be determined). Surrounding the phylogeny are example absorbance spectra that are consistent with the presence of carotenoids; dashed gray lines represent the lower (378nm) and upper (512nm) bounds of carotenoid wavelengths.

### Genotypic diversity

Of our 157 isolates, we successfully obtained sequencing data from 147 (Table S2). Of the 147 successful samples, the average number of Nanopore FASTQ reads that passed our filters was 1,616.8 (± 111 [SE]), with a minimum of 244 reads and a maximum of 6,435 reads.

Our 147 isolates are representative of four phyla, with the two most common being Bacillota (50.3%) and Pseudomonadota (42.8%). These were further subdivided into five classes, with the two most common being Bacilli (50.3%) and Gammaproteobacteria (41.5%). Our isolates belong to 12 orders (the most common being Lactobacillales [43.5%] and Enterobacterales [38.7%]) that encompass 22 families. The most common families include Enterococcaceae and Streptococcaceae (both representing ∼20% of isolates), Yersiniaceae (14.8%), and Erwiniaceae (12.9%). Our samples encompass 35 different genera, with the most common being *Enterococcus* (21.1%), *Lactococcus* (19.7%), *Serratia* (12.2%), and *Pantoea* (10.2%). Genera that were unique to aerobic conditions (n = 21) include *Agrobacterium*, *Bacillus*, *Brucella*, *Delftia*, *Enterobacter*, *Exiguobacterium*, *Kluyvera*, *Kosakonia*, *Kurthia*, *Lysinibacillus*, *Mammaliicoccus*, *Micrococcus*, *Morganella*, *Orbus*, *Plantibacter*, *Priestia*, *Pseudomonas*, *Rothia*, *Staphylococcus*, *Stenotrophomonas*, and *Tsukamurella*. Isolates that belong to the genus *Pantoea* were almost exclusively grown under aerobic conditions, but one isolate grew under anaerobic conditions. Genera that were unique to anaerobic conditions (n = 4) included *Cedecea*, *Providencia*, *Raoultella*, and *Streptococcus*. Genera that grew under both aerobic and anaerobic conditions (n = 10) included *Apibacter*, *Carnobacterium*, *Enterococcus*, *Erwinia*, *Hafnia-Obesumbacterium*, *Klebsiella*, *Lactococcus*, *Pantoea*, *Rahnella*, and *Serratia*. No *Sphingomonas* spp. were recovered on either selective or non-selective media.

All host species except Chestnut-sided Warbler had at least one unique bacterial genus. These included *Agrobacterium*, *Brucella*, *Klebsiella*, *Kurthia*, *Providencia*, *Pseudomonas*, *Rahnella*, *Raoultella*, *Staphylococcus*, *Streptococcus*, and *Tsukamurella*, which were unique to Purple Martins; *Exiguobacterium*, *Mammaliicoccus*, *Plantibacter*, and *Priestia* were unique to Blue-winged Warblers; *Cedecea* and *Hafnia-Obesumbacterium* were unique to Gray Catbirds; *Delftia* was unique to Yellow Warblers; *Micrococcus* was unique to Common Yellowthroats; *Orbus* was unique to Hooded Warblers; and *Kluyvera*, *Kosakonia*, and *Lysinibacillus* were unique to Common Wall Lizards.

We further subdivided the 35 genera into 96 unique ASVs. Most genera were placed into one or two ASVs and three were divided into more than ten ASVs (*Enterococcus* = 19 ASVs; *Pantoea* = 13 ASVs; *Serratia* = 12 ASVs). Most ASVs were unique to a single host species (92.7%). No ASVs were shared by all eight host species, but ASV1 (*Lactococcus*) was shared by six, including Blue-winged Warbler, Common Yellowthroat, Chestnut-sided Warbler, Hooded Warbler, Yellow Warbler, and Purple Martin.

Our phylogeny was highly supported, primarily at deeper nodes, but shallower nodes also had high bootstrap and SH-aLRT support (Figure 2, Figure S2). According to SILVA taxonomy, several genera were non-monophyletic. For example, *Enterococcus* and *Serratia* were polyphyletic, such that their most recent common ancestor encompasses isolates assigned to multiple other genera. *Enterobacter* and *Pantoea*, though they each formed a largely monophyletic clade, exclude one descendent lineage, indicating instances of paraphyly.

PERMANVOA tests indicated that neither host species (R^2^ = 0.2, p = 0.14) nor sampling site (R^2^ = 0.1, p = 0.40) explained any variation between microbiomes (Figure 3). Phylogenetic diversity differed significantly across species (F = 4.33, p = 0.03)—warblers and Purple Martins tended to have higher values than Gray Catbirds and Common Wall Lizards—but did not differ among sampling sites (F = 3.04, p = 0.09; Figure 4).

**Figure 3.**
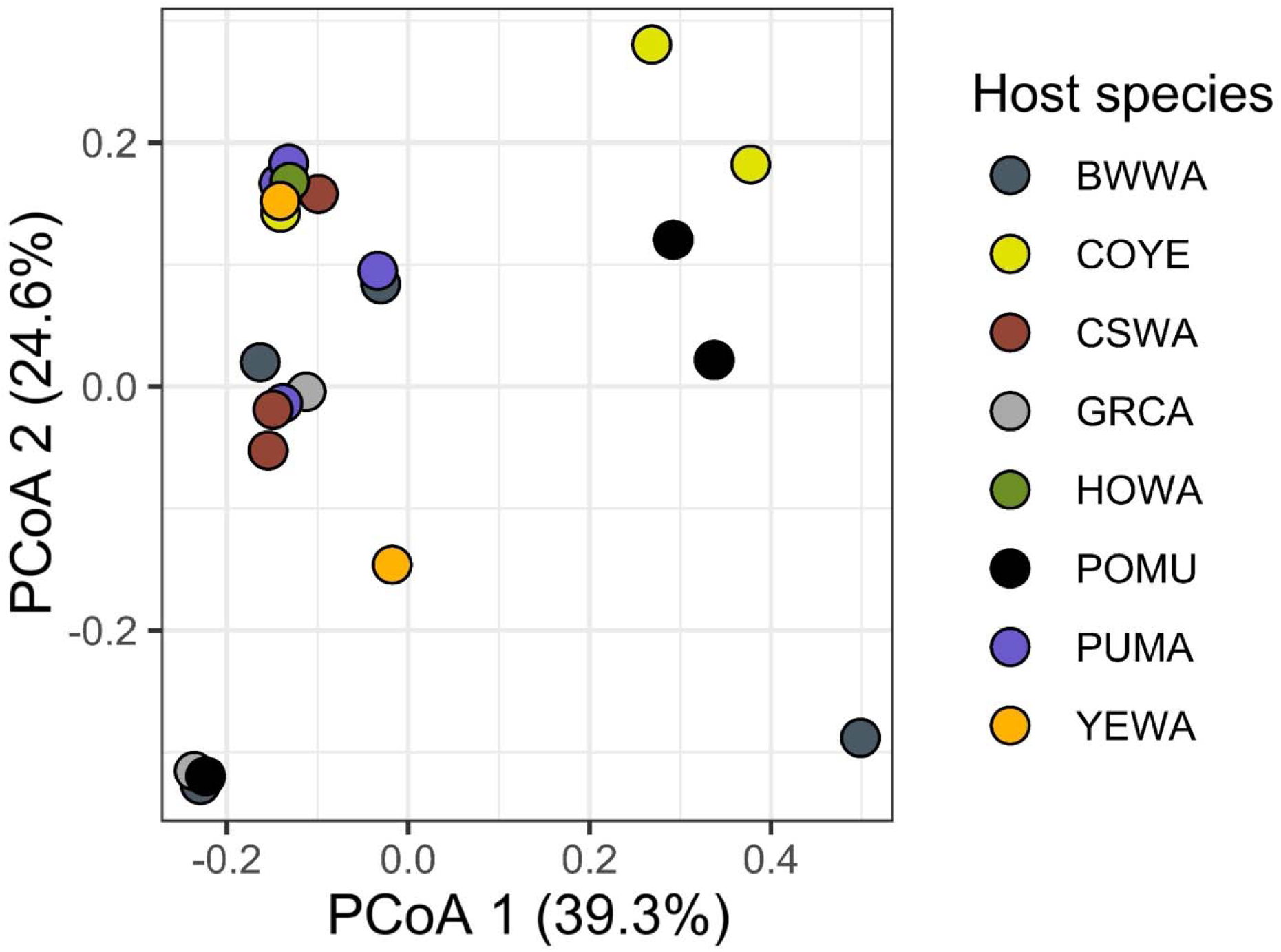
Principal coordinates analysis (PCoA) of bacterial isolates among samples based on ASV presence/absence using unweighted UniFrac distances. Each point represents a sample, and colors indicate host species by their species code.

**Figure 4.**
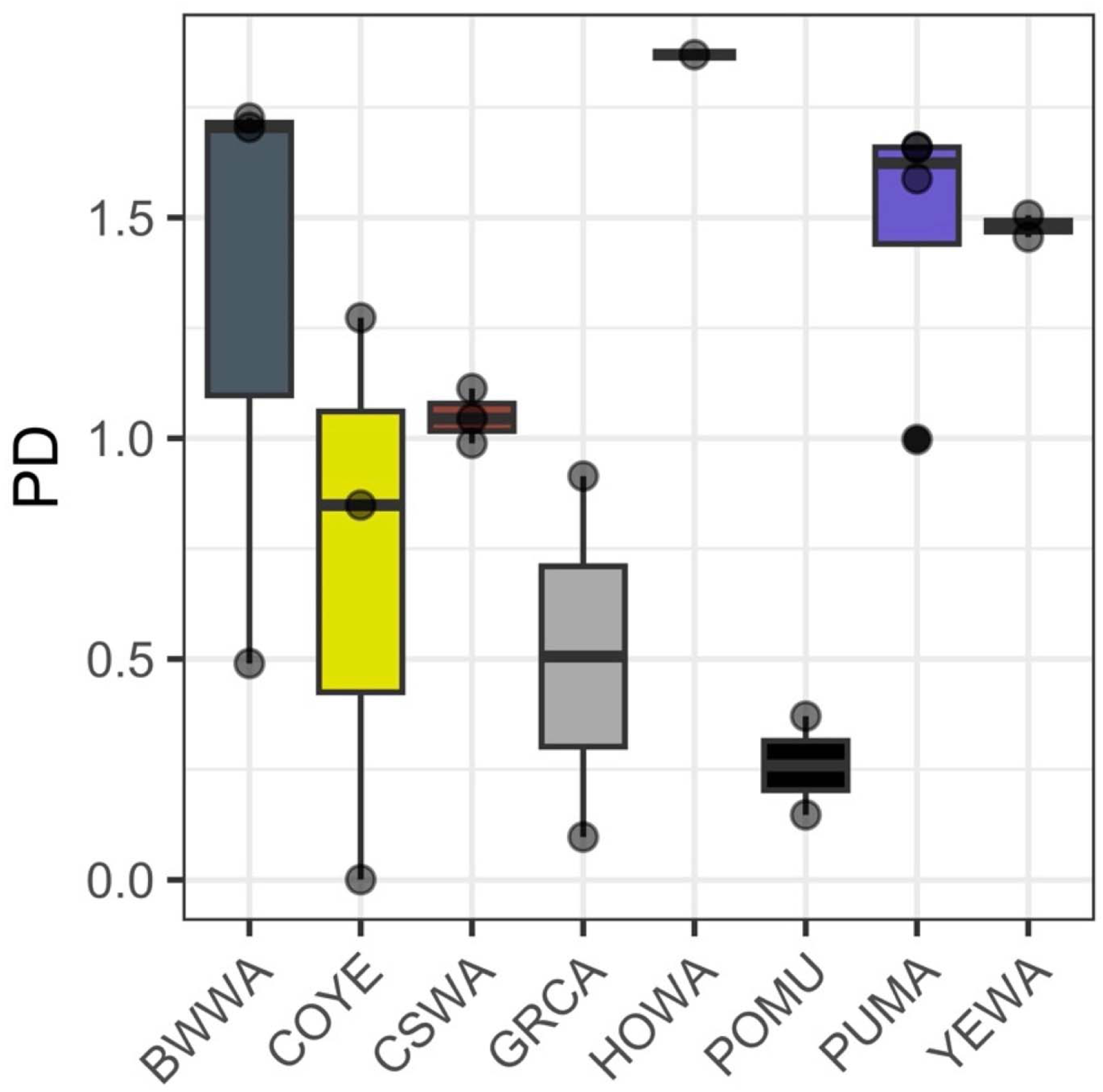
Faith’s phylogenetic diversity (PD) alpha index across host species. Each point represents an individual host and host species are indicated by their species code and associated color.

## DISCUSSION

Our results provide the first direct evidence that gut-associated bacteria isolated from wild sauropsids exhibit diverse pigment phenotypes that are consistent with carotenoids. Broad sampling and high-resolution sequencing demonstrate that these carotenoid-producing bacteria are not restricted to a single taxonomic group, sampling site, or host species (or host family or class), which suggests that they may represent a ubiquitous, yet hidden, component of the sauropsid gut microbiome. Together, these findings provide support for the view that microbial symbionts may be a valuable source of carotenoid pigments to their host.

Although the majority of the isolates were non-pigmented, we frequently observed yellow- or orange-pigmented isolates. Every host species had at least one pigmented isolate, and these phenotypes spanned all four bacterial phyla represented in our dataset. This widespread occurrence across distantly related bacterial and host lineages suggests that carotenoid-like pigments are a recurrent trait in gut-associated bacteria rather than a taxonomic or host-specific anomaly. The yellow phenotype was most often associated with *Pantoea*, which was found only in our avian samples. All but one *Pantoea* were yellow pigmented and exhibited lambda max values within the expected range for carotenoids (Figure 2). *Pantoea* is known to synthesize carotenoids such as zeaxanthin (Misawa *et al*. 1990; Hundle *et al*. 1994), which suggests that the colorful pigments we observed likely reflect genuine functional potential for carotenoid biosynthesis. This interpretation is consistent with the presence of multiple carotenoid biosynthesis genes (e.g., *crt* genes) in a *Pantoea* genome assembly from these samples (RefSeq GCF_054512635.1). Notably, *Pantoea* is also among the most prevalent taxa recovered from 16S metabarcoding across a broad sampling of warbler gut microbiomes (Baiz *et al*. 2023), and was prevalent among isolates from gray-breasted parakeet cloacal swabs (Beleza *et al*. 2021). Thus, despite our (avian) cultures relying on only two media types under two oxygen conditions (an approach that underrepresents unculturable or fastidious taxa), we were still able to isolate some of the most dominant bacterial taxa that are known to be associated with the avian gut microbiome. Altogether, this indicates that our culture-based sampling captured at least a subset (albeit an underrepresentation) of ecologically important and naturally abundant carotenoid-producing bacteria from sauropsid gut microbiomes.

In contrast to the relatively tight correlation between phenotype and genotype observed in *Pantoea*, *Enterococcus* and *Serratia* exhibited more heterogeneity under both aerobic and anaerobic conditions. These two genera were among the most prevalent taxa in our sampling (at least of avian hosts), which is consistent with a previous study of warbler microbiomes which found *Enterococcus* and an unidentified genus within the family Yersiniaceae (of which *Serratia* is a member) as the two most prevalent taxa (Baiz *et al*. 2023). *Enterococcus* was the most ASV-rich genus in our dataset (19 ASVs across 31 isolates), and *Serratia* was among the top three most ASV-rich (12 ASVs across 18 isolates; *Pantoea* exhibited 13 ASVs across 15 isolates). Both *Enterococcus* and *Serratia* produced isolates that fell within and outside of the expected carotenoid lambda max range and encompassed a spectrum of pigmented and non-pigmented color phenotypes (Figure 2). This pattern may suggest considerable genomic variation in the prevalence of *crt* genes in these genera, a trend documented in other bacterial taxa. For example, the distribution of *crtN* and *crtM* genes (which encode for C30 carotenoid biosynthesis) has been found to be widely scattered across taxa within the bacterial family Lactobacillaceae and may be related to differential habitat adaptations (Eilers *et al*. 2024). An alternate possibility may be related to context-specific metabolic expression patterns: because isolates were grown under both aerobic and anaerobic conditions, shifts in pigment production may reflect facultative physiological responses (Pedraz, Blanco Cabra and Torrents 2020), as have been documented in *Enterococcus* (Hagi, Kobayashi and Nomura 2014) and *Serratia* (Rodríguez *et al*. 2023). Targeted genomic screening for *crt* loci and controlled experiments manipulating oxygen availability in these taxa would help to disentangle these possible mechanisms. Taken together, these results highlight that carotenoid-associated traits can vary widely even within bacterial genera, emphasizing the need to consider both genomic and environmental sources of functional diversity in host-associated microbiota.

The confirmed presence of carotenoid-producing bacteria isolated from a diversity of sauropsid hosts adds novel context to the growing number of studies that have found correlations between host microbiomes and variation in phenotype (Nguyen *et al*. 2020; Marques Silva *et al*. 2023; Baiz, Wood and Toews 2024; Koellsch, Poulin and Salloum 2024; Slevin *et al*. 2025). More specifically, these findings suggest that microbial pigment production in the gut may represent a previously unreported contributor to host coloration. Accordingly, three of the four genera most frequently represented in our isolates (*Pantoea*, *Enterococcus*, and *Serratia*) are also among the genera most commonly detected on wild bird feathers (Giorgio *et al*. 2018; Silva *et al*. 2022; Tran *et al*. 2022). The taxonomic overlap between internal gut-associated microbes and external integument-associated microbes supports the possibility of a functional link between carotenoid-producing bacteria in the gut and carotenoid signatures detected in the integument, either directly or indirectly. However, direct comparisons of carotenoids produced by gut-associated bacteria and those deposited in host integument (e.g., feathers, scales) are still lacking. Despite this, predictive metagenomics analyses revealed that carotenoid-associated plumage scores in *Vermivora* (Parulidae) hybrids were correlated with the predicted abundance of KEGG orthologs and enzymes in carotenoid biosynthesis pathways (Baiz, Wood and Toews 2024). More broadly, the ecological or evolutionary mechanisms that enable carotenoid-producing bacteria (and generally any microbe) to persist in the host gut, and the conditions under which they might influence host traits, remain largely unknown (Decaestecker *et al*. 2024).

We observed a similar prevalence of carotenoid-synthesizing bacteria in both avian and lizard hosts (approximately 40%), vertebrate lineages that diverged ∼280 million years ago (Kumar *et al*. 2017). This may suggest a deep evolutionary history of symbiosis between carotenoid-synthesizing bacteria and sauropsid hosts. Several factors may explain the prevalence of carotenoid-synthesizing bacteria in the sauropsid gut microbiome. First, because carotenoids protect bacteria from environmental stressors, they tend to flexibly inhabit and thrive under various conditions without strict niche specialization (Martino *et al*. 2016; Eilers *et al*. 2024). Thus, they may be rather ubiquitous across host habitats and, coincidentally, disperse to and colonize the gut microbiome frequently. Consistent with this view, we isolated carotenoid-synthesizing bacteria from every host species sampled at each of our study sites, and neither site nor host species were strong predictors of microbiome composition (Figure 3). Similarly, many of these taxa may be common symbionts of prey, like *Pantoea* which is commonly associated with insects (Walterson and Stavrinides 2015), providing a plausible route of acquisition and explain its ubiquity among our samples. Alternatively, and regardless of their baseline environmental prevalence), hosts may selectively filter for carotenoid-synthesizing bacteria. Filtering could occur indirectly by some physiological mechanism and allow for the proliferation of carotenoid-synthesizing taxa over others (e.g., gut pH, temperature) or more directly if natural selection favors hosts that harbor high abundances of carotenoid-synthesizing bacteria. Carotenoids are a highly valuable resource for animals (Blount 2004; Fiedor and Burda 2014; Toews, Hofmeister and Taylor 2017), and if hosts are able to take advantage of carotenoids synthesized by gut microbes in addition to dietary sources, we would expect selection to favor the maintenance of such a mutualism. However, carotenoid expression may be condition dependent (Hagi, Kobayashi and Nomura 2014), and further research is necessary to determine whether the bacteria we isolated here synthesize carotenoids within the host gut (i.e., not just in cultured isolation) and, if so, what functional role they may play.

Despite their potential importance, virtually nothing has been documented about the prevalence or diversity of carotenoid-producing gut bacteria in wild hosts. Our results suggest that some of the most prevalent bacteria in wild sauropsid gut microbiomes are capable of producing carotenoids and are taxonomically widespread. These findings raise the possibility that the host gut microbiome serves as a previously underappreciated reservoir of biologically relevant and valuable carotenoids, although future work is needed to determine the identity of these compounds (e.g., using high-performance liquid chromatography [HPLC]). Furthermore, understanding the extent to which hosts may exploit microbially derived carotenoids in their own physiological processes remains an important target for future research. More broadly, this work contributes to a growing body of evidence that microbial symbionts can play hidden roles in shaping host traits with potential consequences for the ecology and evolution of their hosts.

## ACKNOWLEDGEMENTS

We thank Asela Wijeratne, Kipa Tamrakar, and the T. Krabbenhoft Lab for helping with PCR and Nanopore sequencing design; Sarah Walker, Trevor Krabbenhoft, and Zhen Wang for access to lab equipment; Heather Williams, Shawn Healy, Tina Nguyen, and Erin Karnatz for assistance in the field; Samantha Fontaine for assistance with developing culturing protocols, absorbance assays, and lizard collections in the field; and Allison Remick and Isabella Harter for assistance with lizard microbial culturing. Birds were captured and handled under the United States Geologic Survey Bird Banding Laboratory Permit #24043 and the University at Buffalo IACUC #PROTO202400015. Lizards were captured and handled under the Ohio Division of Wildlife Scientific Collection License (#SC230080) and Kent State University IACUC (Protocol 560 BT 24-02).

---

Figure S1. Absorbance spectra for all isolates; dashed gray lines represent the lower (378nm) and upper (512nm) bounds of carotenoid wavelengths.

Figure S2. Maximum likelihood phylogenetic tree estimated from the 16S rRNA gene of 147 bacterial isolates cultured from wild sauropsid fecal samples. Tips are labeled with the genus and ASV of each isolate. Nodes with strong support (SH-aLRT ≥ 80% and ultrafast bootstrap ≥ 95%) are denoted by yellow circles. The scale bar represents the number of nucleotide substitutions per site.

**Table S1.**
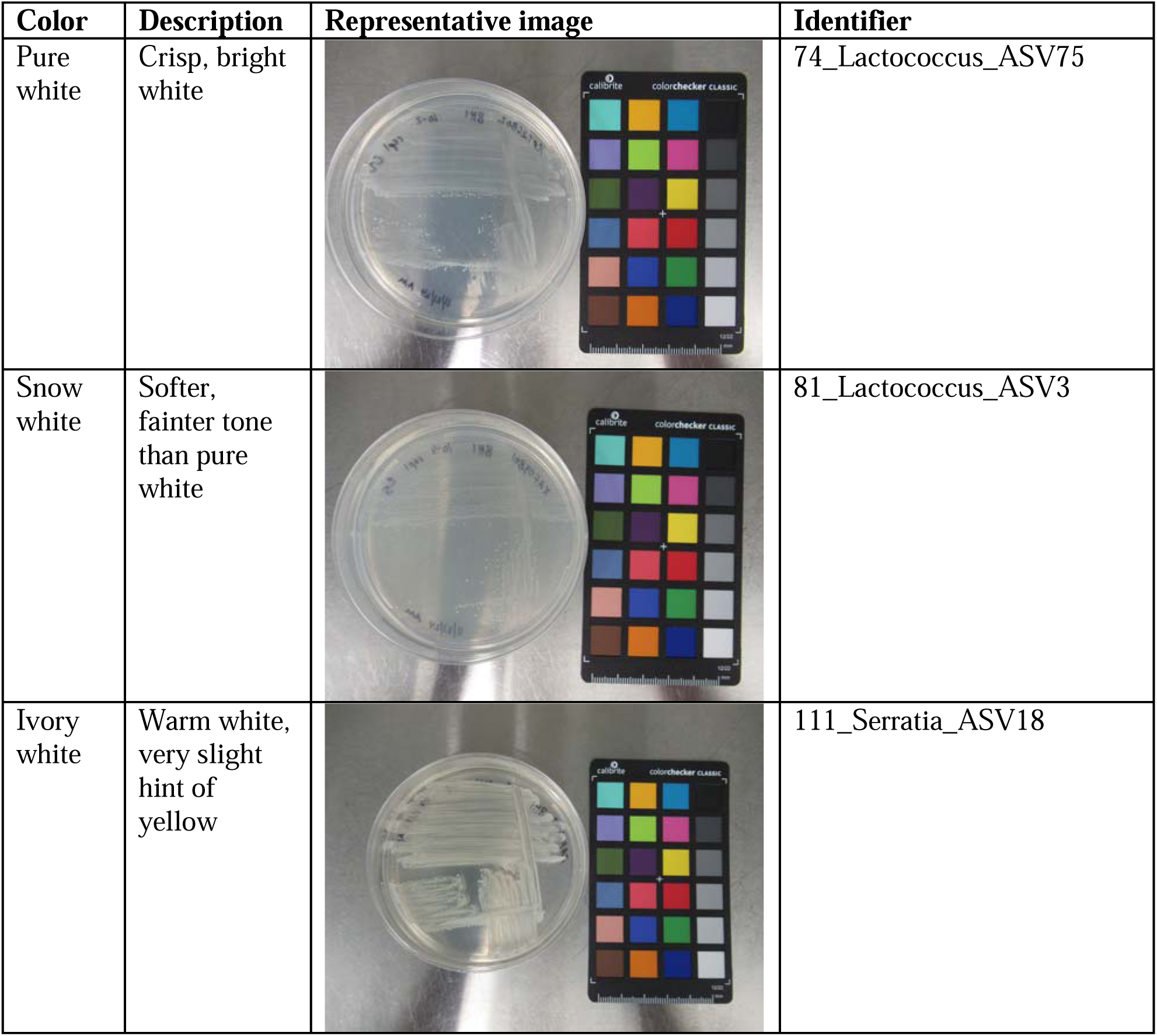
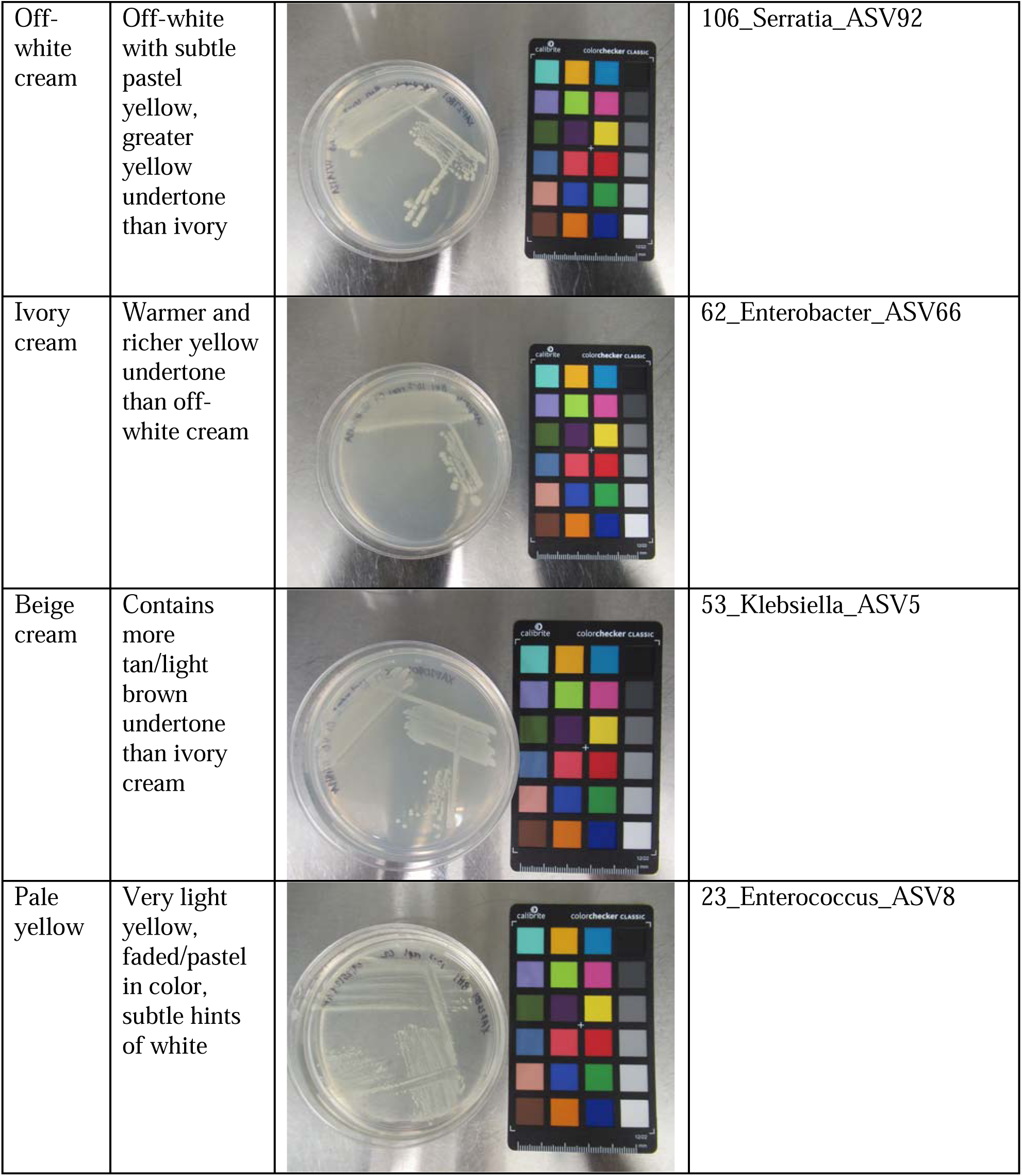
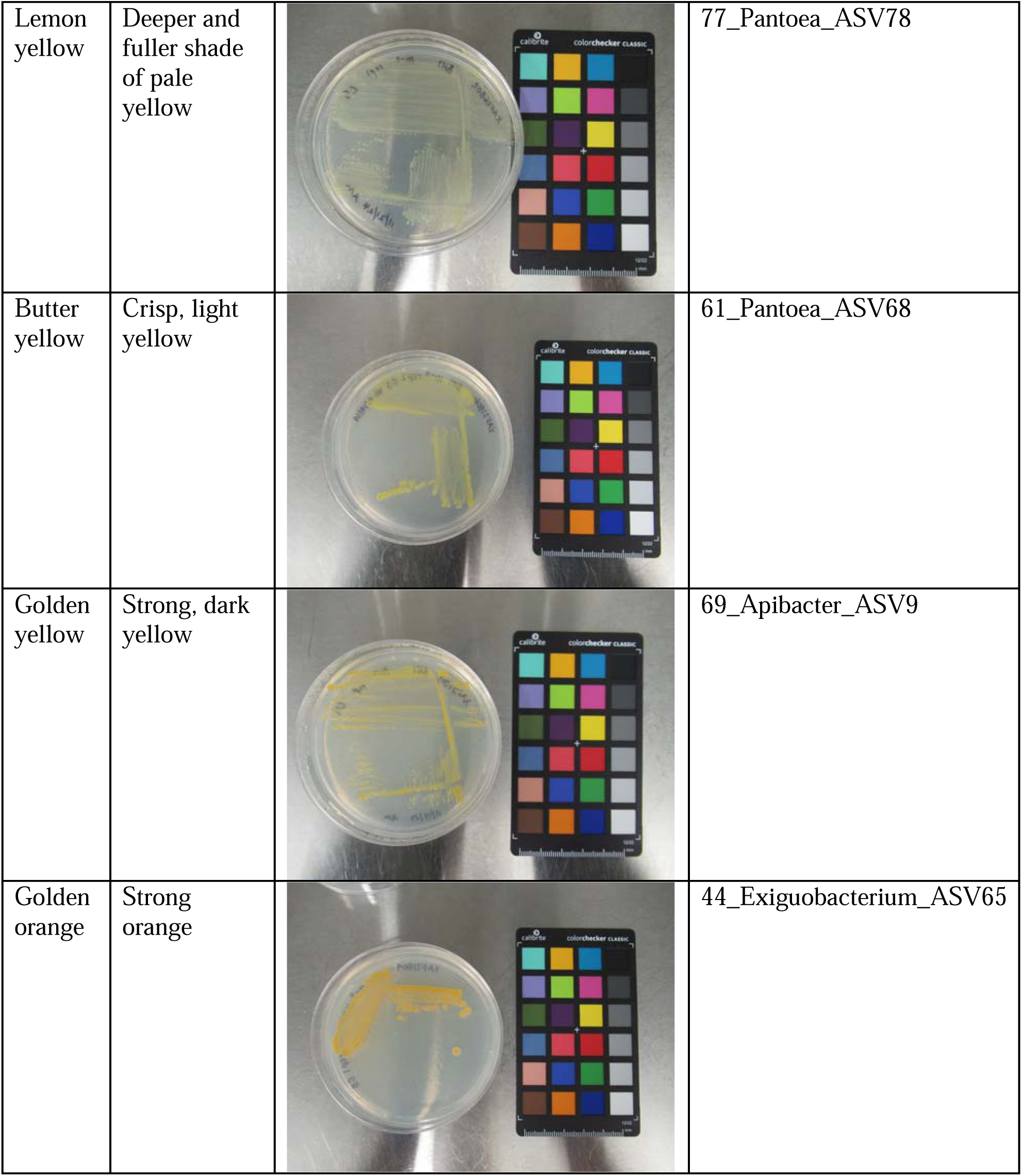
Summary of phenotype-based color categories. Each row includes the color name, a description of each color’s distinguishing characteristics, a representative image of an isolate for each color, and the individual identity that links the representative isolate to the labels on the phylogeny and absorbance assay curves.

| Color | Description | Representative image | Identifier |
| --- | --- | --- | --- |
| Pure white | Crisp, bright white |  | 74_Lactococcus_ASV75 |
| Snow white | Softer, fainter tone than pure white |  | 81_Lactococcus_ASV3 |
| Ivory white | Warm white, very slight hint of yellow |  | 111_Serratia_ASV18 |
| Off-white cream | Off-white with subtle pastel yellow, greater yellow undertone than ivory |  | 106_Serratia_ASV92 |
| Ivory cream | Warmer and richer yellow undertone than off-white cream |  | 62_Enterobacter_ASV66 |
| Beige cream | Contains more tan/light brown undertone than ivory cream |  | 53_Klebsiella_ASV5 |
| Pale yellow | Very light yellow, faded/pastel in color, subtle hints of white |  | 23_Enterococcus_ASV8 |
| Lemon yellow | Deeper and fuller shade of pale yellow |  | 77_Pantoea_ASV78 |
| Butter yellow | Crisp, light yellow |  | 61_Pantoea_ASV68 |
| Golden yellow | Strong, dark yellow |  | 69_Apibacter_ASV9 |
| Golden orange | Strong orange |  | 44_Exiguobacterium_ASV65 |

Table S2. Metadata associated with each isolate including: host species code (host_species), host species scientific name (host_species_sci), the individual host record identifier (host_record_id), the collection date formatted as dd-mmm-yyyy (collection_date), collection site name (site), country and state of collection site (geo_loc_name), the isolate identification number (isolate_id), isolate phenotype data (color, form, surface_texture, margin, opacity, elevation, size), the type of media the isolate was cultured on (media_type), whether the isolate was cultured under aerobic or anaerobic conditions (culture_type), taxonomy of the isolate (phylum, class, order, family, genus), sequencing technology for the isolate (seq_type), the position on 96-well plates for absorbance assays (curve_plate_position), phylogeny label with OTU identifier (phylogeny_curve_otuid), phylogeny label with ASV identifier (phylogeny_curve_asvid), a simplified ID number (clean_id_number), the OTU identity (otu_id), the ASV identity (asv_id), whether the absorbance curve was consistent with a carotenoid (carot_profile), and the NCBI record numbers (SRA_or_GenBank).

