## Supplementary figures and images for "Hidden palette: Culturing the sauropsid gut microbiome reveals a high prevalence and diversity of bacteria that undergo carotenoid biosynthesis"

### Figure S2

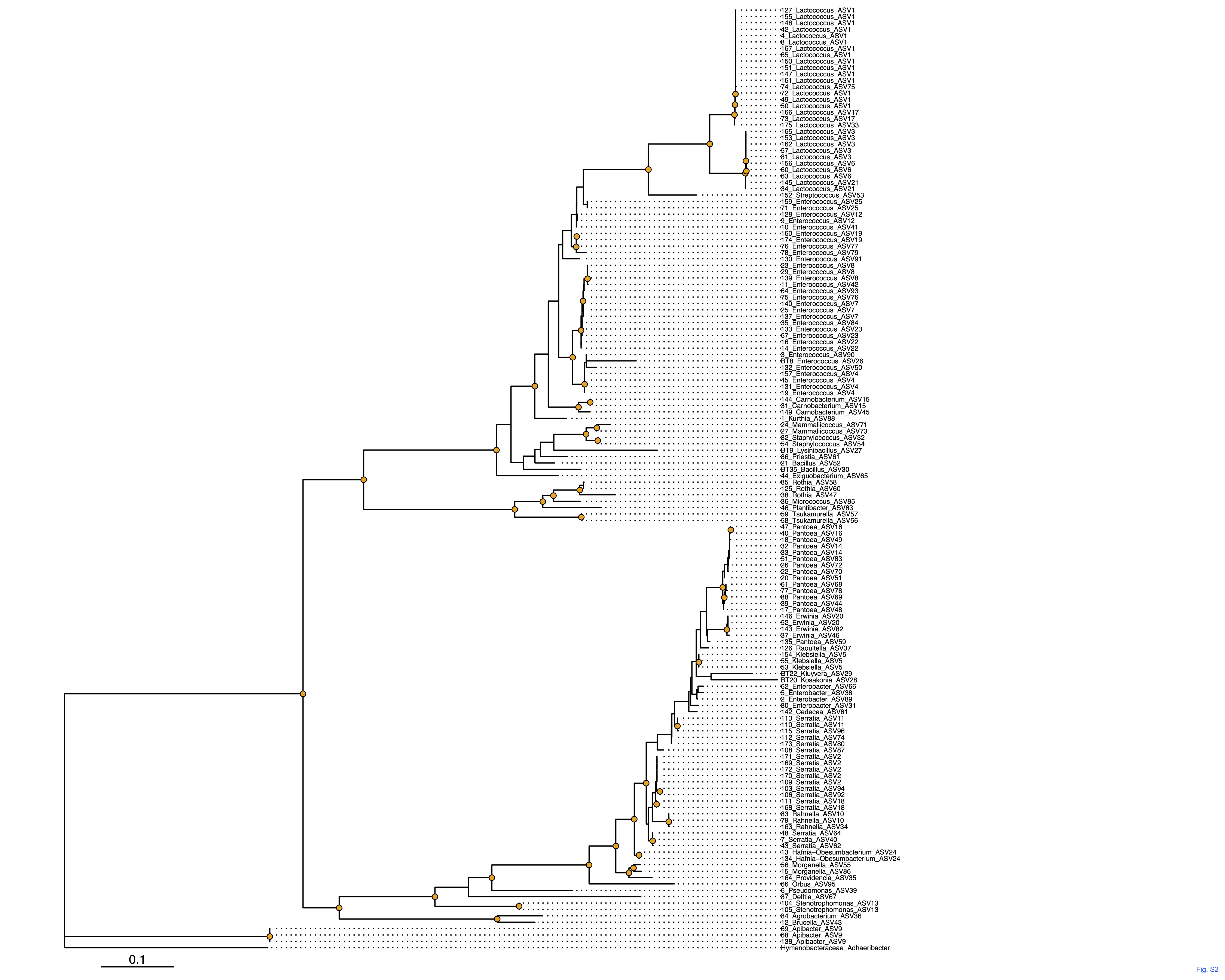
